# Early Visual Cortex Supports One-Shot Episodic Memory via Spatially Tuned Reactivation

**DOI:** 10.1101/2025.04.04.647327

**Authors:** Robert Woodry, Jonathan Winawer, Serra E. Favila

## Abstract

Episodic memory retrieval is thought to rely on the reactivation of prior perceptual representations in sensory cortex, a phenomenon known as cortical reinstatement. Support for this idea in early sensory areas comes from memory studies involving repeated exposure and explicit recall instructions. In everyday life, however, people rely on details from single events without repeated practice or instruction. We used fMRI to test whether memory responses emerge in early visual cortex after a single encoding event. Twenty adults viewed objects presented once in one of four parafoveal locations, then later completed object recognition (old/new) and location recall tasks in which objects were shown foveally. Both memory tasks evoked spatially tuned, albeit substantially lower amplitude, responses compared to encoding in V1 to V3. These responses were tuned to the encoded parafoveal location, even though the recognition task did not ask the participant to remember the encoded location. These responses were more robust and precise for successfully remembered items, linking neural tuning in early visual cortex to one-shot memory performance. Our findings demonstrate that early visual cortex supports the defining property of episodic memory, the ability to retrieve sensory details from a single event.

**Significance Statement:** Episodic memory allows us to vividly recall details from single past experiences. While it is hypothesized that this “mental time travel” relies on reactivating sensory cortex, previous studies used highly practiced associations or explicit visualization cues, which differ from how we naturally recall unique events. In this fMRI study, participants viewed a stream of unique objects at parafoveal locations. We found that later retrieval of these objects spontaneously reactivated early visual cortex in a spatially specific manner even when spatial location was not probed: The memory reactivated the same retinotopic location where the object initially appeared. Furthermore, the precision of this reactivation predicted memory success. These findings demonstrate that sensory cortex is fundamental to the rapid, one-shot learning that characterizes human episodic memory.

## Introduction

The defining feature of episodic memory is the ability to retrieve the sensory details of a single, once-experienced event. Dominant models propose that the hippocampus rapidly encodes new experiences and then later retrieves them by reinstating encoding-related cortical activity patterns (Damasio, 1989; McClelland et al., 1995; Schacter et al., 1998). In this view, cortical reinstatement is the mechanism by which episodic memory recovers perceptual detail and guides action.

Human neuroimaging and neurophysiology studies provide compelling evidence that episodic memory retrieval reinstates high-level visual features such as item category (Kuhl et al., 2011; Kuhl & Chun, 2014; Minxha et al., 2020; Polyn et al., 2005; Wadia et al., 2024). However, category-level information cannot guide action; the brain must also recover low-level spatial coordinates. Space acts as a high-resolution scaffold, binding items into coherent, actionable events. While the spatial circuitry in early visual cortex is well-suited for representing spatial information (Barlow, 1986), and its contribution to working memory is an active area of inquiry (Dake & Curtis, 2025), its crucial role in supporting episodic memory remains underexplored.

While some long-term memory studies report spatial reinstatement in primary visual cortex, they rely on highly-trained cue-stimulus associations and explicit instructions to recall visual details (Breedlove et al., 2020; Favila et al., 2022; Favila & Aly, 2023; Vo et al., 2022; Woodry et al., 2025). Though useful for reliably activating the cortex, visualization instructions can engage different mechanisms than those used for memory (Farah, 1984; Rosenbaum et al., 2004; Siena & Simons, 2024)(Corkin, 2002; Zeman et al., 2015) and repetitive associative learning alters memory content (Bridge & Paller, 2012). Previous studies thus do not test the neural mechanisms that retrieve details from a single event, an ability that is fundamental to episodic memory. If reinstatement in early visual cortex reflects the mechanism by which episodic memory accesses spatial information, it should emerge whenever a single event is remembered. It therefore remains unclear whether spatial reinstatement in early visual cortex is a core feature of episodic memory.

Memory for a single event is not simply a weaker version of memory for a highly learned association. Repetition and retrieval practice strengthen recall (Roediger & Butler, 2011; Young & Bellezza, 1982), but they also alter how memories are represented and retrieved. With repetition, memory representations can become more dependent on cortex (Frankland & Bontempi, 2005) and more gist-like, as they are built up by averaging over many related experiences (McClelland et al., 1995). Although changes to memory representations can happen with the passage of time even without repeated exposure, repeated study and retrieval have been proposed to accelerate this process (Antony et al., 2017; W. Yu et al., 2024). As a result, reinstatement observed after extensive training may reflect representations that emerge through repetition rather than those engaged when remembering a single, unique event. This distinction between retrieval of single events and of repeated associations is particularly relevant for early sensory cortex, where involvement in higher-level cognition remains debated (Linton et al., 2024) and evidence for reactivation has been less consistent than in higher-level areas (Kosslyn & Thompson, 2003).

Here, we directly tested whether remembering a once-experienced event re-engages spatial representations in early visual cortex. To test this prediction, we scanned 20 participants while they performed a one-shot memory task involving parafoveal encoding of objects, followed by foveal old/new recognition, and foveal spatial recall. We used population receptive field (pRF) encoding models (Dumoulin & Wandell, 2008) to quantify cortical reinstatement during recognition and recall. Traditional univariate analyses and decoding approaches would likely fail to capture subtle, low-amplitude one-shot memory signals. Our use of encoding models therefore provides a sensitive test of whether the human episodic memory system can use the high-resolution architecture of the early visual cortex to reconstruct the past. Finally, by linking spatial reactivation strength to trial-by-trial memory performance, we evaluate whether early visual cortex recruitment is a functional requirement for the successful retrieval of a unique episode.

## Results

We scanned 20 subjects while testing their memory for parafoveally viewed objects across three interleaved tasks (Figure 1). During *encoding*, they viewed trial-unique objects at one of four polar angles, two degrees away from fixation. During *recognition*, they viewed objects foveally and made old vs. new judgments. During *recall*, they viewed old objects foveally and reported the location at encoding. We measured spatial responses in early visual cortex during all three tasks. All claims reported in the main text also hold after excluding the two experimenter subjects (Supplemental Table 1).

**Figure 1.**
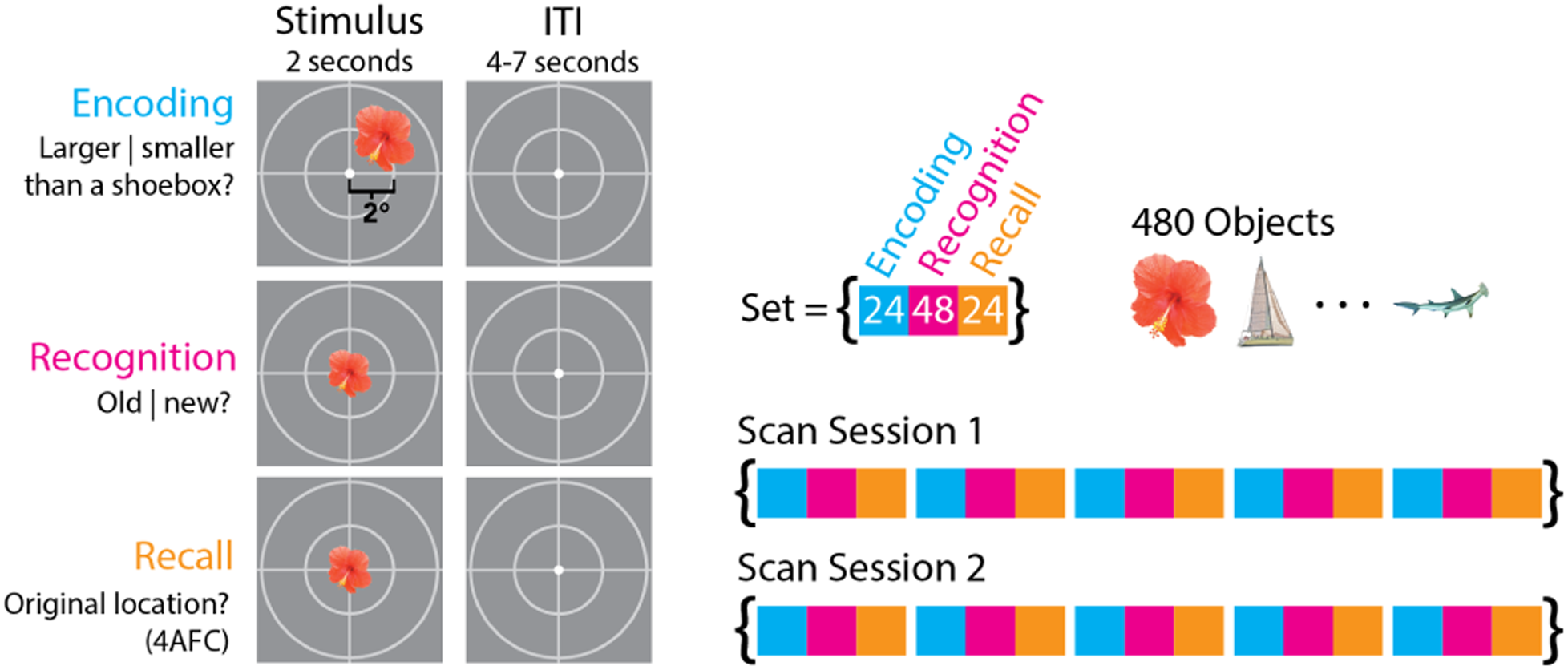
Experimental design. Subjects participated in two fMRI sessions with 15 scans each: encoding, recognition, and recall, repeated five times. The encoding and recall scans included 24 trials each, corresponding to 24 distinct objects, in random order. Recognition scans contained the same 24 objects, with an additional 24 new objects (lures), shown in random order. In encoding trials, subjects were briefly shown an object in one of four locations at 2° eccentricity (45°, 135°, 225°, or 315° polar angle) and asked to indicate whether they were larger or smaller than a shoebox in real life. In recognition trials, subjects were shown an object at fixation and were asked to indicate whether it was old or new. In recall trials, subjects were shown an old object at fixation, and were asked to indicate its original location.

### Memory performance for object recognition and spatial location recall

We first quantified performance on the recognition and recall tasks. We designed the experiments so that participants would be above chance in both tasks but well below ceiling in the recall task, enabling us to assess trial-to-trial variability in behavior and brain responses.

Performance was good in both tasks. In the recognition task, the hit rate was 90% [95 CI 87%, 92%] and the false alarm rate was 7% [5%, 10%] (Figure 2A). In the recall task, the average accuracy was 69% [95 CI 63%, 74%], with a chance level of 25% (Figure 2B). Performance was stable across scans except for a slight improvement in the recall task after the first block (Figure 2C, D).

**Figure 2.**
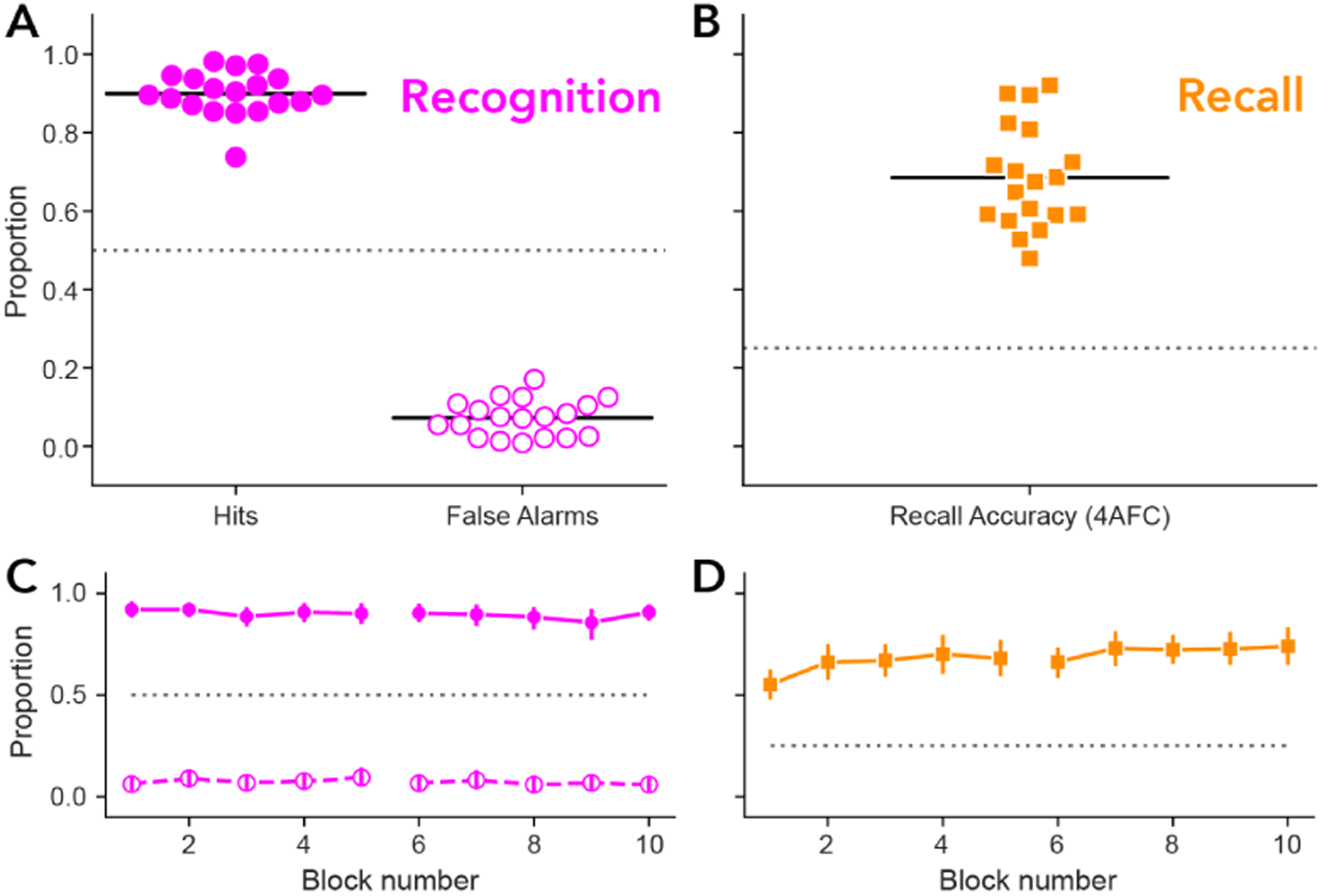
Behavioral performance on recognition and recall tasks. A) Subject-level performance on the recognition task. Individual subjects are plotted as circles for hit rate (filled) and false alarm rate (unfilled). Black horizontal lines indicate means across subjects. The dotted line indicates chance performance. B) Subject-level performance on the recall task. C) Recognition performance across blocks. Data points are the means across subjects for hit rate (filled) and false alarms (unfilled). Error bars indicate 95% CI obtained from 10,000 bootstraps across subjects. D) Recall performance across blocks. The code and data to generate this and all subsequent figures can be found at https://osf.io/a9hkg/.

### A single viewing is sufficient for spatially tuned memory responses in early visual cortex

To assess how retrieving these object memories affected activity in visual cortex, we projected the activity from the surface of early visual cortex (V1-V3) to visual space using pRF model parameters (Figure 3A). From these projections we computed polar angle activation profiles for each visual map and task, and quantified each profile’s spatial tuning by its peak location, amplitude, tuning width, and fidelity.

**Figure 3.**
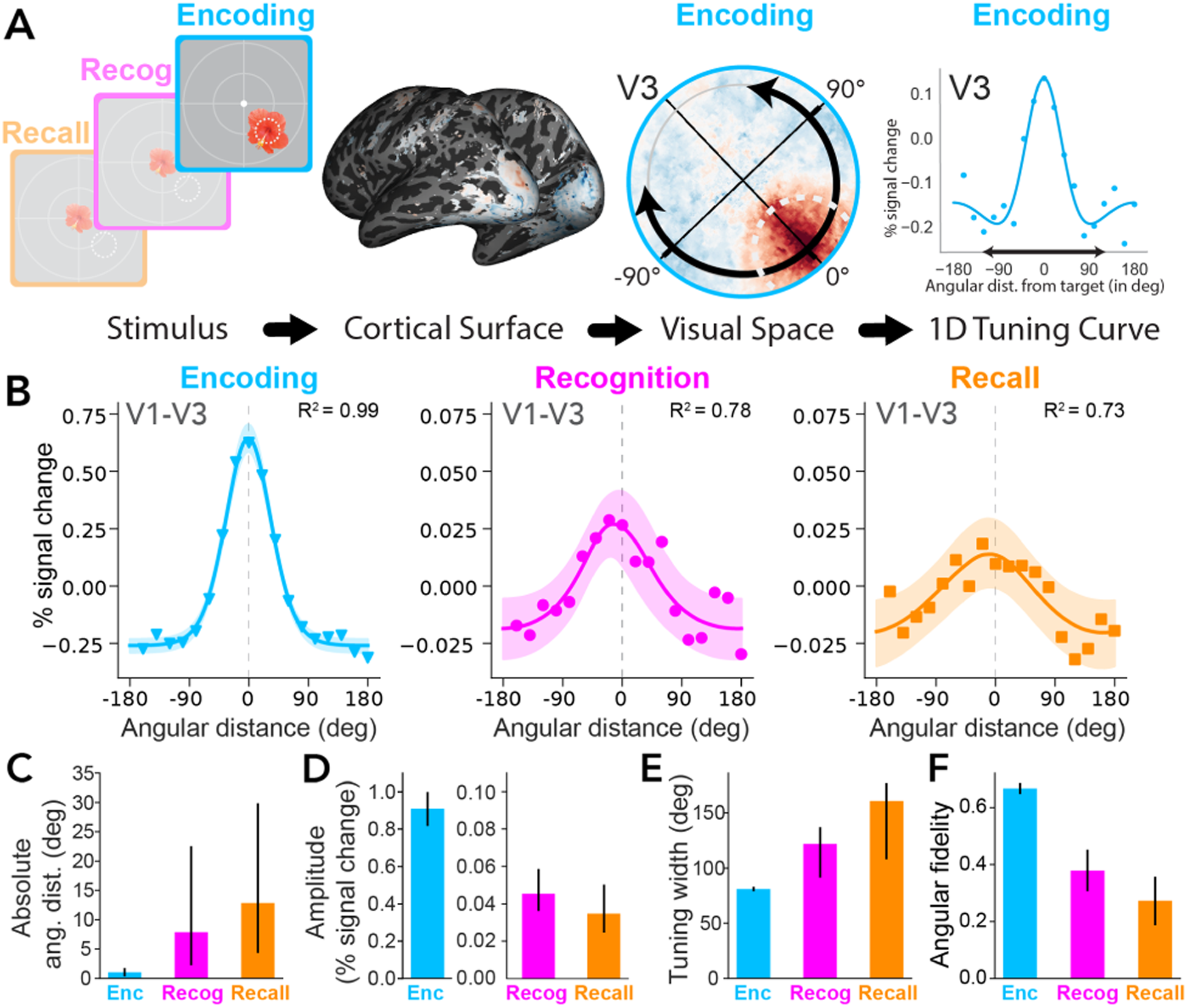
Single-shot memory responses in visual cortex are tuned to encoded location. A) Mapping BOLD responses to polar angle activation profiles. A stimulus evokes a BOLD response in visual cortex. This activity is projected to visual space and binned by angular distance from target, producing a polar angle activation profile. This example profile is from V3 during the encoding task. B) Polar angle activation profiles in V1-V3 across the three tasks. Shaded regions indicate 68% confidence intervals across subjects. C-F) Estimates of tuning quality for the three tasks. Error bars indicate 68% confidence intervals from bootstrapping across subjects. For data plotted separately by visual area (V1, V2, V3) or by individual participant, see Supplemental Figure 3-1 and Figure 4, respectively. C) Distance between peak location and object source location, computed as absolute deviation of the peak of the polar angle activation profile. D) Response amplitudes (peak to trough). The y-axes have different scales for encoding and retrieval. E) Tuning width (full-width at half the maximum amplitude). F) Angular fidelity, a summary metric that depends on the peak location, width, and signal-to-noise ratio in the response profiles

**Supplemental Figure 3-1.**
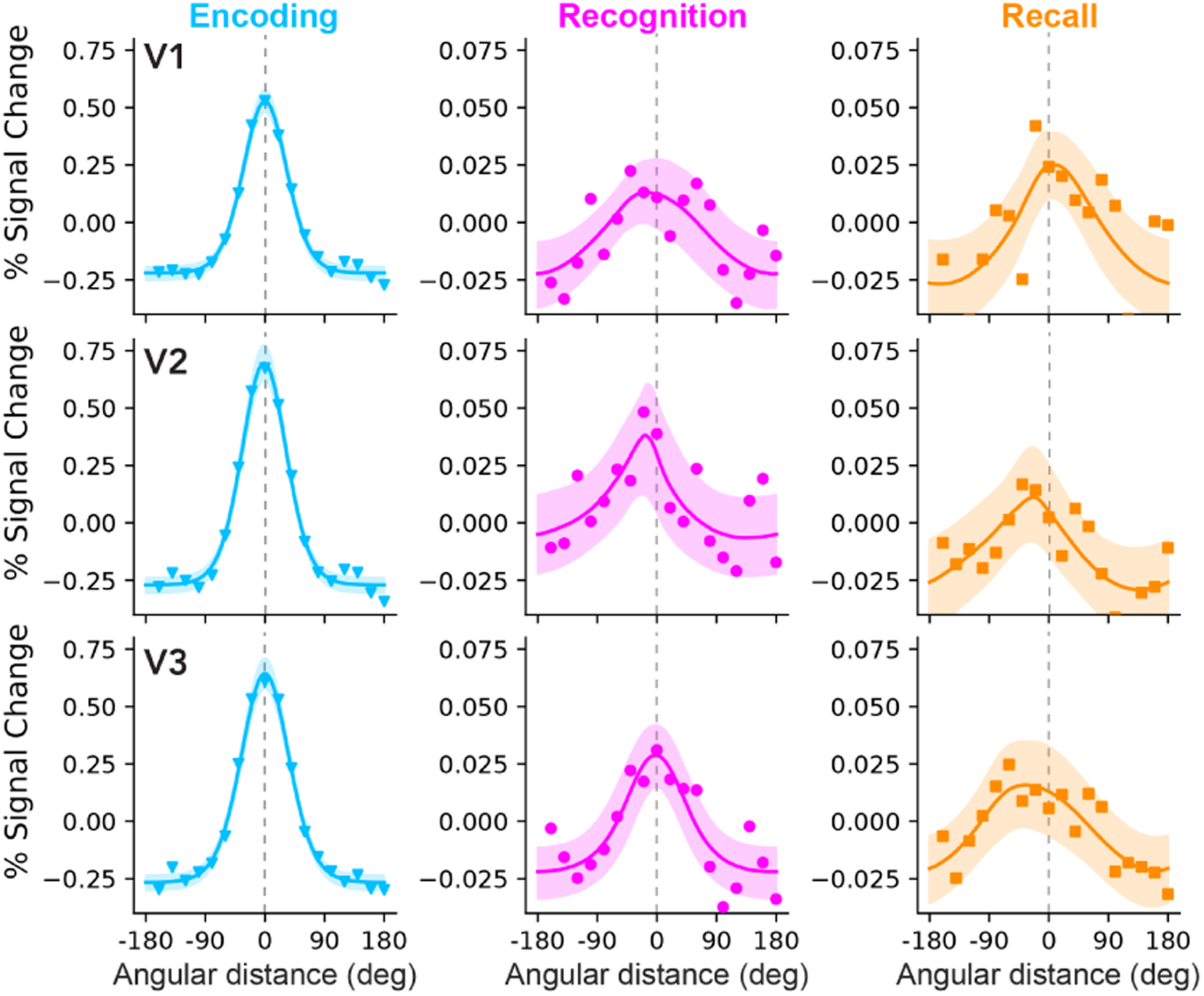
Spatial tuning during encoding, recognition and recall for V1, V2, and V3. Polar angle activation profiles during encoding, recognition, and recall for each visual map V1, V2, and V3. Spatial tuning is evident within each visual map. V1 encoding fidelity was at 0.65 [95% CI 0.58, 0.74], recognition was 0.31 [0.1, 0.42], and recall was 0.26 [0.13, 0.33]. V2 encoding fidelity was at 0.63 [0.6, 0.68], recognition was 0.27 [-0.05, 0.48], and recall was 0.17 [-0.05, 0.4]. V3 encoding fidelity was at 0.71 [0.63, 0.73], recognition was 0.35 [0.14, 0.5], and recall was 0.32 [0.1, 0.41].

For each task, the center of the polar angle activation profiles was close to the target location, indicating that activity in V1-V3 was tuned to the object’s angular location at encoding. The average distance between the activation profile and the stimulus location during encoding trials was 1° [95% CI 0°, 2°], during recognition 8° [0°, 31°] and during recall 13° [0°, 48°] (Figure 3B; see visual areas plotted separately in Supplemental Figure 3-1). The activation profiles for the encoding data were well fit by the von Mises functions, with R^2^ of 0.99. The recognition and recall data were noisier, but the von Mises functions still accounted for over 70% of the variance in the activation profiles. Tuned responses during the recognition task is especially notable as location was not probed for this task. Moreover, the tuning was evident from the first recognition memory block in the experiment, before participants had a chance to learn that they would also be tested on location memory (Supplemental Figure 3-2).

**Supplemental Figure 3-2.**
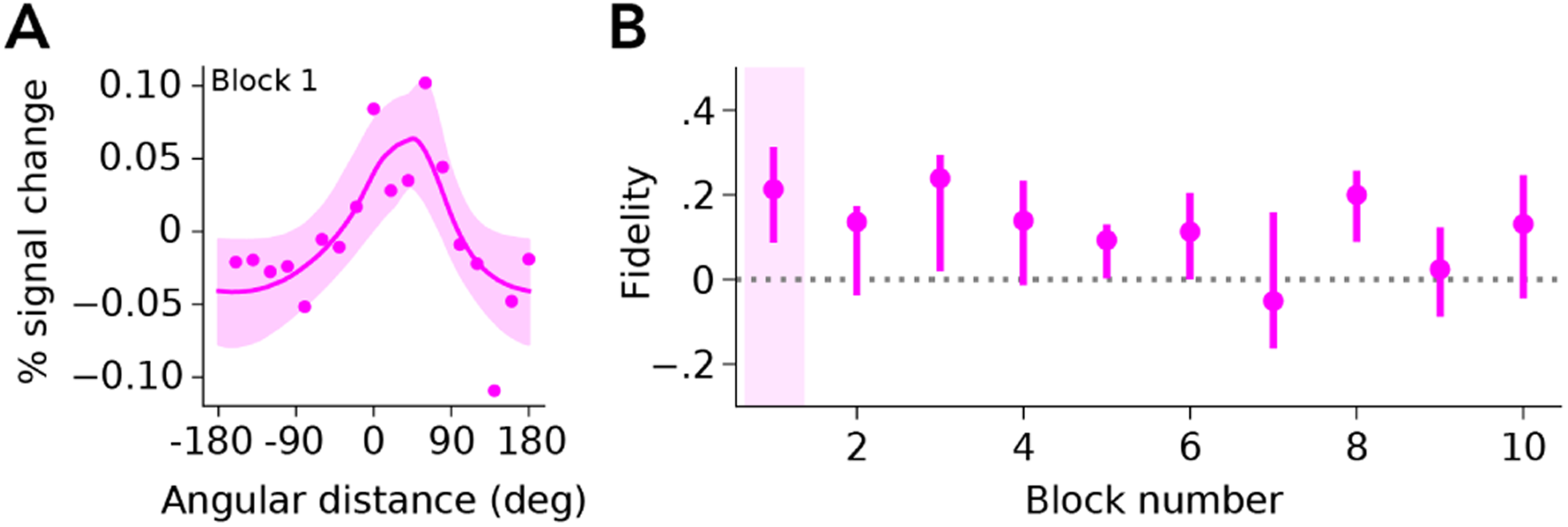
Recognition memory response by block number in V1. A) Polar angle activation profile for V1 during the first recognition block, restricted to trials with subsequent accurate recall (i.e. location) memory. The first recognition block differs from the others in that participants are naive to the recall task in the first recognition block. Even so, the recognition memory response showed robust tuning near the original object’s location, with a fidelity of 0.21 [95 CI -0.02, 0.42]. B) Fidelity of the recognition memory response across blocks. Shaded region denotes block 1 plotted in panel A. Error bars are 68% confidence intervals bootstrapped across subjects. The fidelity of the recognition memory response does not increase in subsequent blocks, which shows that knowing about the recall task does not improve the recognition memory response.

The activation profiles differed substantially between encoding, recognition, and recall even though all three were centered close to the encoding location. First, the amplitude was much larger for encoding than recognition and recall: the encoding response was 0.86% signal change [95 CI 0.69%, 1.03%] larger than the recognition response, and 0.87% [0.7%, 1.03%] larger than the recall response (Figure 3D). Expressed as a ratio, the recognition and recall response amplitudes were about 25 times lower than the encoding responses (0.9 vs 0.035). Second, the responses during both memory retrieval tasks were more broadly tuned than those at encoding. The width of the activation profile was 81° [68 CI 79°, 83°] for encoding, 122°, [68 CI 97°, 136°] for recognition, and 161° [68 CI 111°, 177°] for recall (Figure 3E). Broader tuning in early visual cortex during memory retrieval is consistent with our previous work (Favila et al., 2022; Woodry et al., 2025). The amplitude of these memory responses, however, is notably lower than in the prior studies.

The combination of greater peak angular distance and broader tuning during memory retrieval compared to encoding is reflected in lower *angular fidelity*. Fidelity quantifies the degree to which the tuning peaks at the target location, ranging from -1 (if the only non-zero response is in the bin 180° away from the target) to +1 (if the only non-zero response is in the bin centered at 0°). A fidelity of 0 indicates no directional bias toward 0° or 180°. This measure is a useful summary of the polar angle activation profile as fidelity is influenced by several factors: fidelity is high when the center position is close to 0°, when the tuning is narrow, and when the data have low noise. The encoding response had a fidelity of 0.67 [95% CI 0.62, 0.71], higher than both memory retrieval tasks: fidelity for recognition was 0.38 [0.11, 0.49] and recall was 0.32 [0.07, 0.46] (Figure 3F). Although the fidelity was lower during recognition and recall than during encoding, it was well above 0, indicating reliable spatial tuning in both memory retrieval tasks.

The spatial tuning to the remembered stimulus during recognition and recall trials was not due to eye movement artifacts. Participants were instructed to fixate centrally throughout the trial. Had they not done so, for example saccading toward the remembered location, the presence of the central object stimulus would have caused a retinotopic activation opposite the remembered location, contaminating our tuning function analyses. This did not happen, however – participants maintained good fixation throughout the experiment and exhibited no systematic bias in small gaze deviations during the memory retrieval tasks (Supplemental Figure 3-3).

**Supplemental Figure 3-3.**
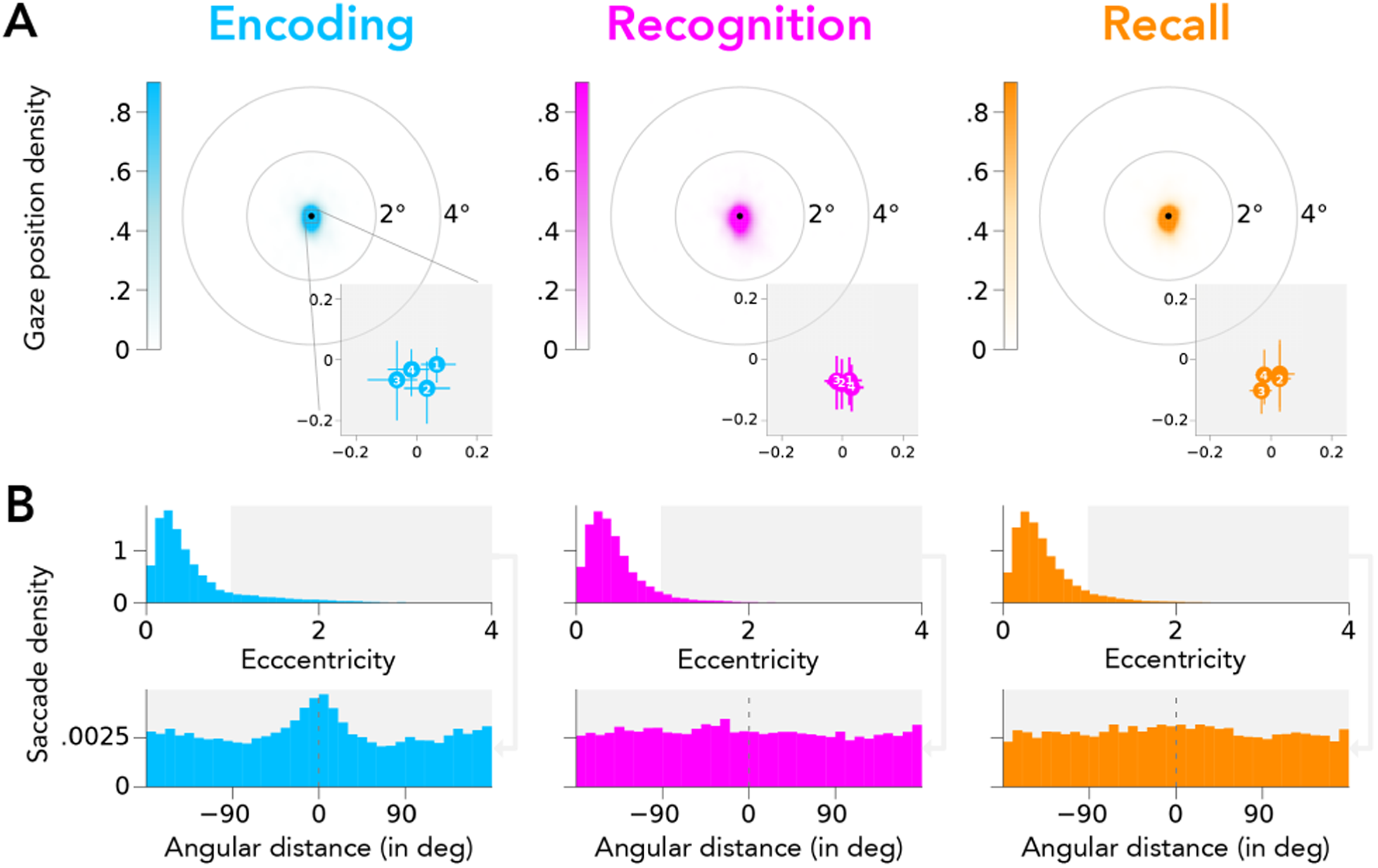
Gaze position and saccades in during each task’s stimulus display. A) Density plot of average gaze position within each trial’s 2-s stimulus display window. Each object’s source location was at 2° eccentricity. Shaded inset: Group-level mean gaze position for different stimulus source locations revealed only minor deviations from fixation point across all tasks (<= 0.1 deg) (n = 13). Error bars represent the 95% confidence intervals of the mean. White numbers indicate source location (1: 45°, 2: -45°, 3: -135°, 4: 135°). The average calibration accuracy of the eye tracker across participants (who had at least two validation runs) was 0.64° [95 CI 0.49°, 1.04°], which is consistent with Eyelink’s reported calibration accuracy levels of 0.5°. B) Top row: Density histogram of saccade eccentricity within each trial’s stimulus display window. Saccades were rare (8.7% of sampled frames across participants [68 CI 7.3%, 10.1%]). Less than 10% of saccades landed closer to the target eccentricity than to the fixation dot (shaded region) in both memory retrieval tasks (encoding = 13.6%, recognition = 8.1%, location = 8.8%). Bottom row: Density histogram of saccade polar angle during stimulus display window for saccades that landed closer to the target eccentricity than fixation point, aligned to the source location of the trial’s object. Dashed gray lines mark polar angle location of stimuli (0°). Aligning near-target saccades to their respective trial’s source location reveals a small bias to the source location during the encoding task, but not during either memory retrieval task. These results demonstrate that subjects are maintaining central fixation throughout the experiment, and that the tuned recognition and recall responses are not an artifact caused by saccades biased towards each object’s source locations.

### Spatially tuned memory responses are evident in individual subjects

We averaged data across participants before computing polar angle response functions for the main analyses above to increase the signal-to-noise ratio, which tends to be low in the memory retrieval tasks. Nonetheless, the overall patterns are evident in most of the individuals (Figure 4). Angular fidelity, computed separately for each participant, was positive in all three tasks: 0.6 [95% CI 0.57, 0.63] in the encoding task, 0.19 [0.1, 0.26] in the recognition task, and 0.15 [0.07, 0.22] in the recall task. For each participant, we compared their measured fidelity to the fidelity derived from a null distribution. The fidelity measure was more than 2 standard deviations greater than the mean of the null distribution for all 19 participants in the encoding condition, 14 of 19 in recognition, and 13 of 19 in recall. Overall, the individual participant analyses indicate that spatial tuning in visual cortex during memory retrieval was not an artifact of averaging across participants, but was instead evident in most individuals.

**Figure 4.**
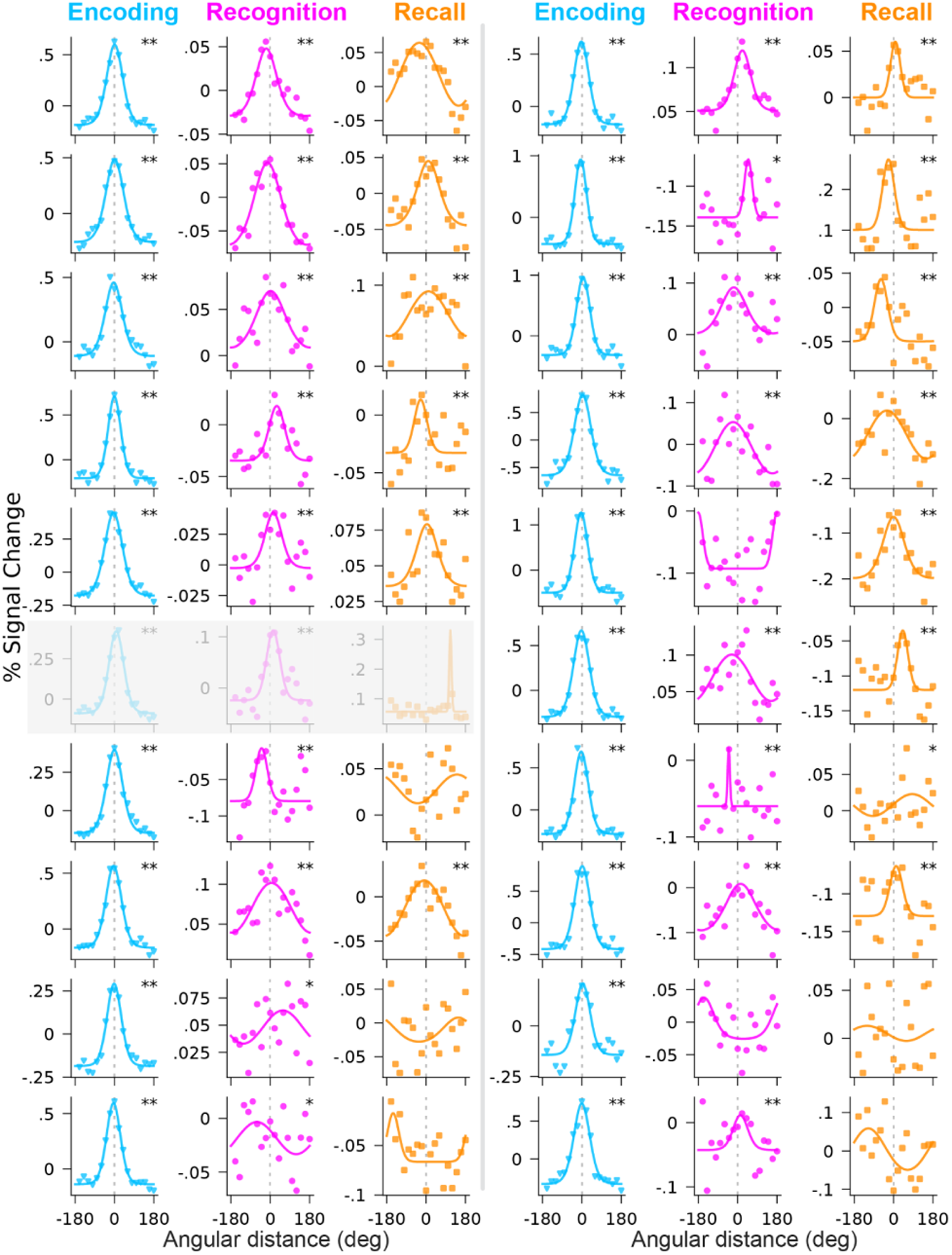
Spatial tuning during encoding, recognition, and recall for individual subjects. V1-V3 polar angle activation profiles for all subjects (n=20) and tasks. Transparent gray overlay indicates the subject excluded from group-level analyses due to not following response report instructions during the recall task. The second and third row in the first column belong to the two experimenter subjects. Dashed grey lines indicate 0°, i.e., the polar angle location of the object at encoding. Asterisks denote responses with angular fidelity greater than 1 (*) or 2 (**) sd from the mean of the null distribution. The null distribution was computed separately for each subject and each condition by scrambling the assignment of target locations across trials, and then re-computing the fidelity, 1,000 times per participant.

### Memory responses in early visual cortex track locations of remembered objects

To test whether spatially tuned memory responses in visual cortex predict successful recall, we grouped trials in each memory retrieval task conditioned on whether the object’s location was later remembered or forgotten (behavioral report during recall, Figure 5A). For each subject, we balanced remembered and forgotten conditions by sub-sampling trials from the condition with more trials to match the number of trials in the condition with fewer trials. We then recomputed the polar angle activation profile for each condition, weighting each subject by the number of trials they contributed to the group average. (Figure 5B). These profiles reveal clear spatial tuning, during both recognition and recall, when the object’s location was successfully remembered. By contrast, this response was less tuned for trials where the object’s location was forgotten, particularly during the recall task. Indeed, the fidelity of the recognition response dropped from 0.35 [95% CI 0.13, 0.44] for remembered trials to 0.24 [-0.08, 0.34] for forgotten trials, while the fidelity of the recall response dropped from 0.35 [0.09, 0.52] to -0.03 [-0.14, 0.18]. The relation between memory performance and fidelity is particularly evident in V1 (Supplemental Figure 5-1). These results reveal that accurate spatial tuning of memory responses in early visual cortex is associated with accurate spatial recall.

**Figure 5.**
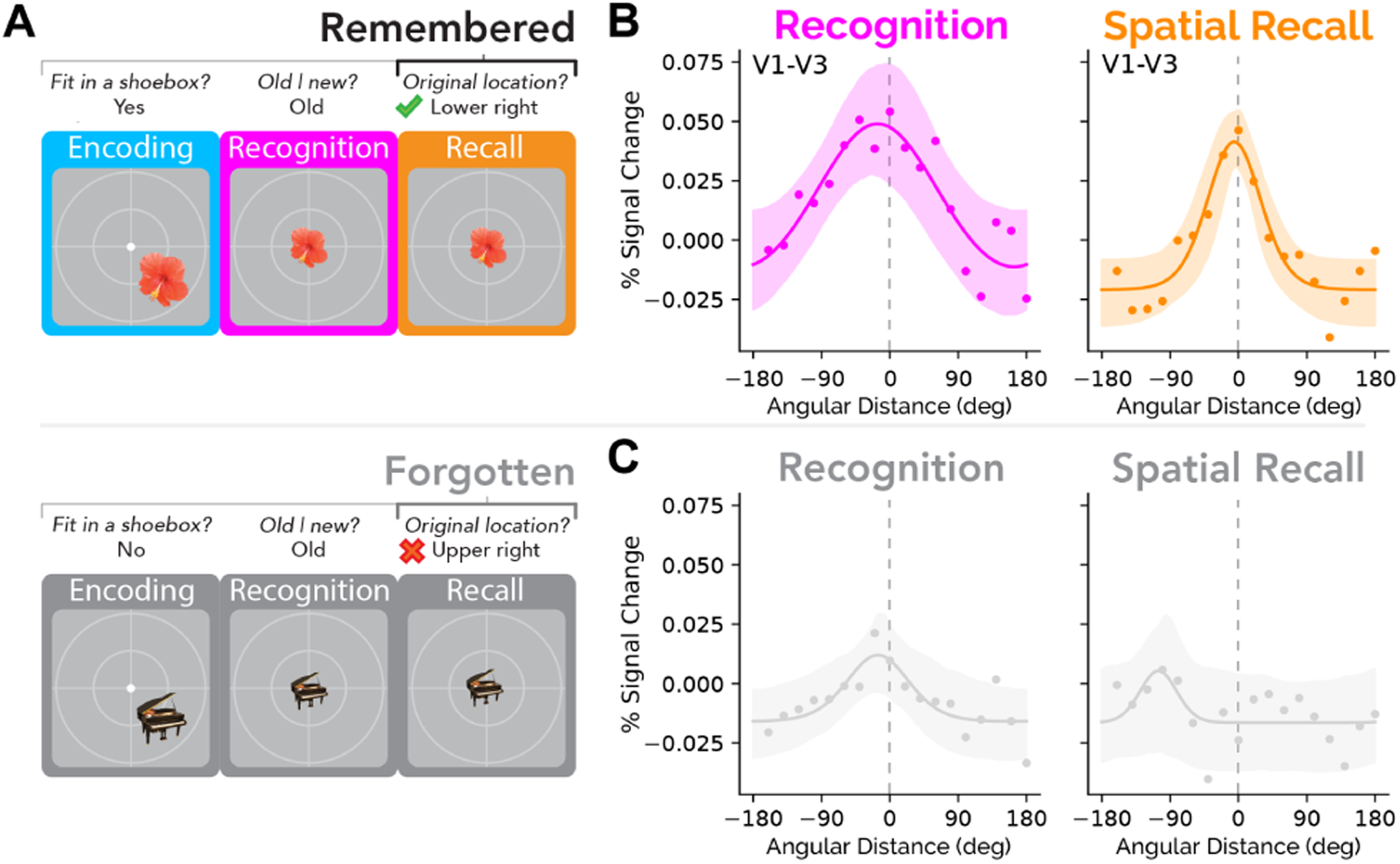
Spatially-tuned memory responses in early visual cortex track recall success. A) Schematic for grouping remembered/forgotten trials across tasks. Trials in each task (encoding, recognition, and recall) are assigned to the ‘remembered’ group if the object’s location is reported correctly during the recall task. Trials are deemed ‘forgotten’ if location is reported incorrectly during the recall task. B) Polar angle activation profiles during both memory retrieval tasks, for trials where the object’s location was remembered at recall. C) Polar angle activation profiles for trials where the object’s location was forgotten at recall.

**Supplemental Figure 5-1.**
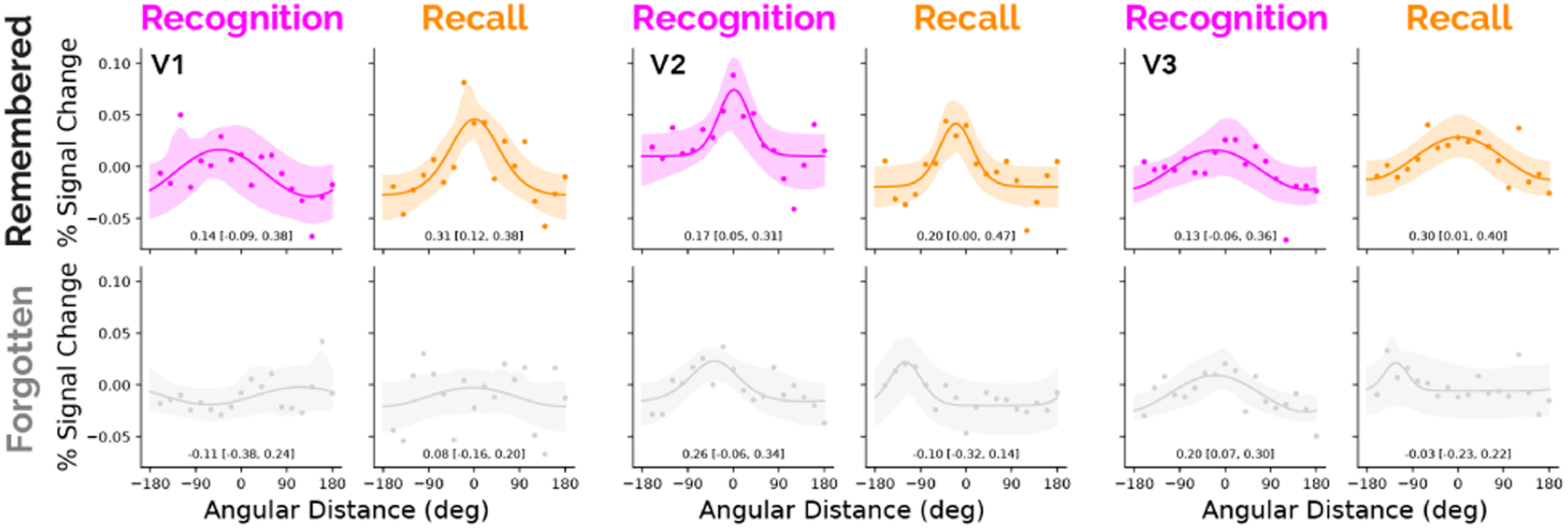
Memory responses for remembered vs. forgotten trials in V1, V2, V3. Polar angle activation profiles for V1, V2, and V3 during both memory retrieval tasks. Trials are grouped according to whether the object location probe was answered correctly (top row) or not (bottom row) during the recall task. Numbers inside the subplots indicate the fidelity of the polar angle activation profile and its 95% confidence interval. The drop in fidelity between remembered and forgotten trials is most evident in V1.

Still, it’s possible that there are spatially tuned responses for forgotten trials, but not at the encoded location. If participants were either retrieving an incorrect memory or guessing and then attending to the reported location, we might expect tuning to the reported location rather than the correct location. To assess this possibility, we repeated the prior analyses but defined 0° as the reported location. The fidelity of the forgotten response remained low across early visual cortex during both recognition and recall: recognition fidelity dropped from 0.35 [95% CI 0.13, 0.44] for remembered trials to 0.25 [-0.13, 0.44] for forgotten trials, and recall fidelity dropped from 0.35 [0.09, 0.52] for remembered trials to 0.11 [-0.21, 0.3] for forgotten trials. This pattern was consistent within each visual map (see Supplemental Figure 5-2). That the report-aligned recall data were not tuned on forgotten trials suggests that the act of reporting the location is not responsible for the tuning during remembered trials.

**Supplemental Figure 5-2.**
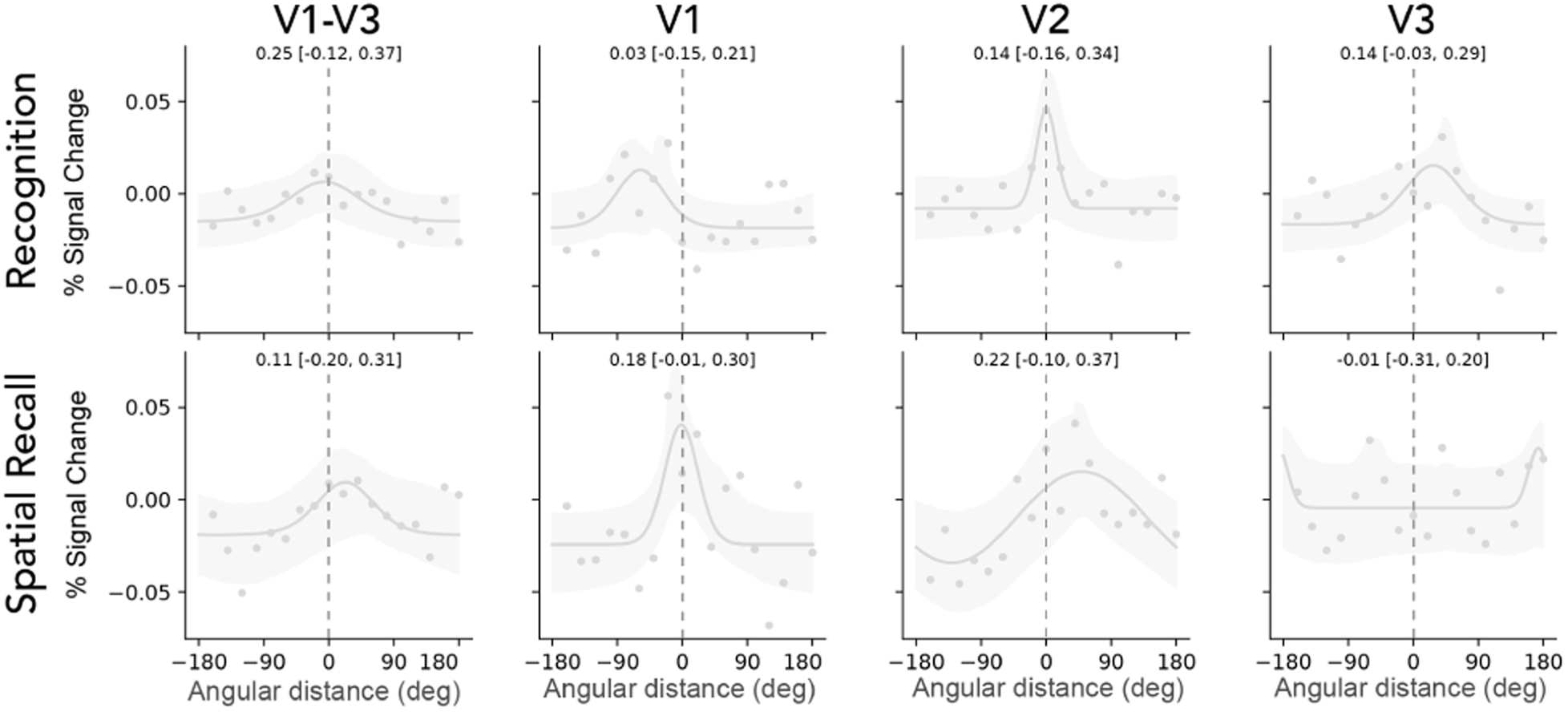
Report-aligned memory responses for forgotten trials for all ROIs. Report-aligned polar angle activation profiles for V1-V3 and each visual map during both memory retrieval tasks for trials where the object’s location was forgotten. Numbers inside the subplots correspond to the fidelity of the polar angle activation profile and its 95% confidence interval. The tuning of the recognition and recall responses for forgotten trials was low across ROIs when the responses were aligned to the behavioral report.

### Spatial tuning at encoding predicts subsequent memory

Objects whose locations were remembered during the recall task may differ in how they were encoded, reflected by differences in cortical activity during stimulus presentation. This is typically referred to as a “subsequent memory effect”. Previous research has generally failed to observe subsequent memory effects in early visual cortex (Kim, 2011). However, prior studies tended to average visual cortex responses across maps, and subsequent memory effects might be limited to subpopulations of visual cortex. Our polar angle activation profiles provide a more precise way to quantify subsequent memory effects.

We tested for subsequent memory effects in visual cortex by comparing the spatial tuning profiles from the encoding task for subsequently remembered vs subsequently forgotten trials (Figure 6A). From these initial comparisons we found slightly larger responses during encoding in each visual area for trials whose object locations were later remembered than for those later forgotten. Specifically, the amplitude (peak minus trough) of the population response for remembered trials was 0.11% BOLD signal change [95 CI 0.01%, 0.22%] larger than the population response for forgotten trials in early visual cortex (Figure 6B). This subsequent memory effect is spatially selective, with the largest difference at the target location (0°), and the smallest difference furthest from the target location (Figure 6C). Because the difference is positive at 0° and negative near 180°, the average difference between the curves is close to 0, which might be why other studies did not find subsequent memory effects in visual cortex.

**Figure 6.**
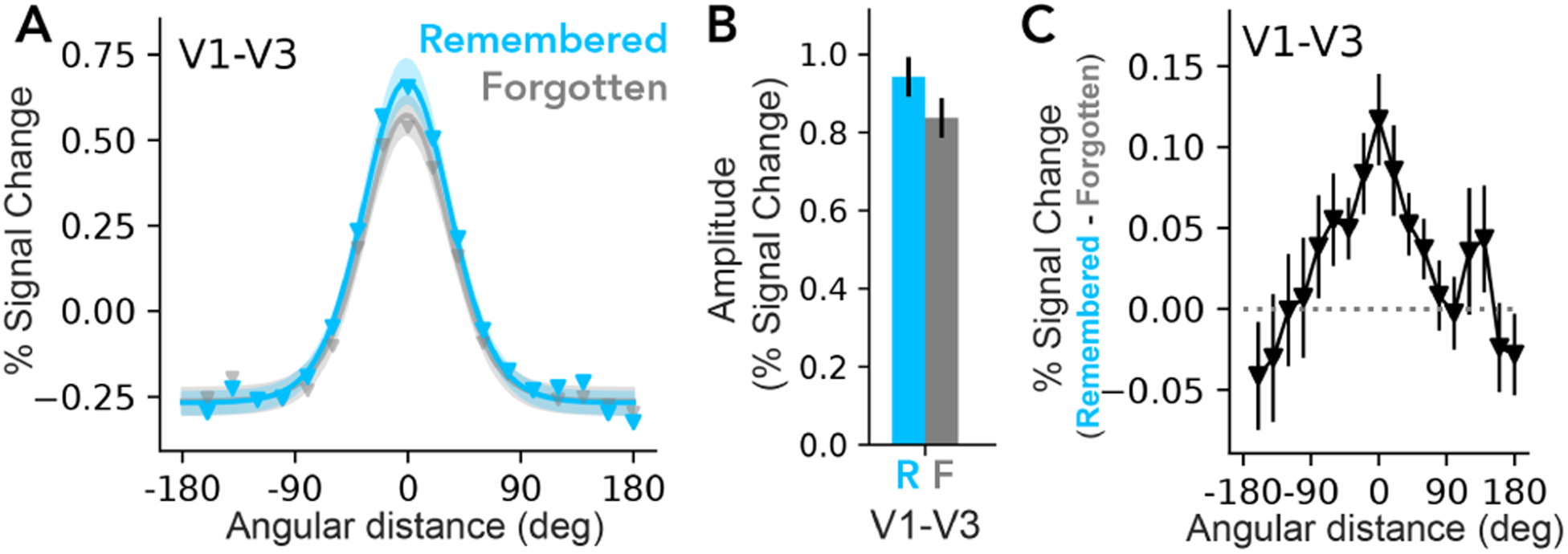
Spatial tuning in early visual cortex at encoding predicts subsequent recall. A) Polar angle activation profiles from early visual cortex during encoding, grouped by whether the encoded object’s location was later remembered (blue) or forgotten (gray) during the recall task. Shaded region indicates 68% confidence interval across subjects. B) Amplitude estimates (peak to trough) during encoding for remembered and forgotten groups. Error bars indicate one standard deviation of the amplitude difference between the two groups across bootstraps. C) Differences in BOLD response between remembered and forgotten groups as a function of angular distance to the target. Error bars indicate one standard deviation of the bootstrapped differences between the two groups. For separate plots of V1, V2, and V3, see Supplemental Figure 6-1.

**Supplemental Figure 6-1.**
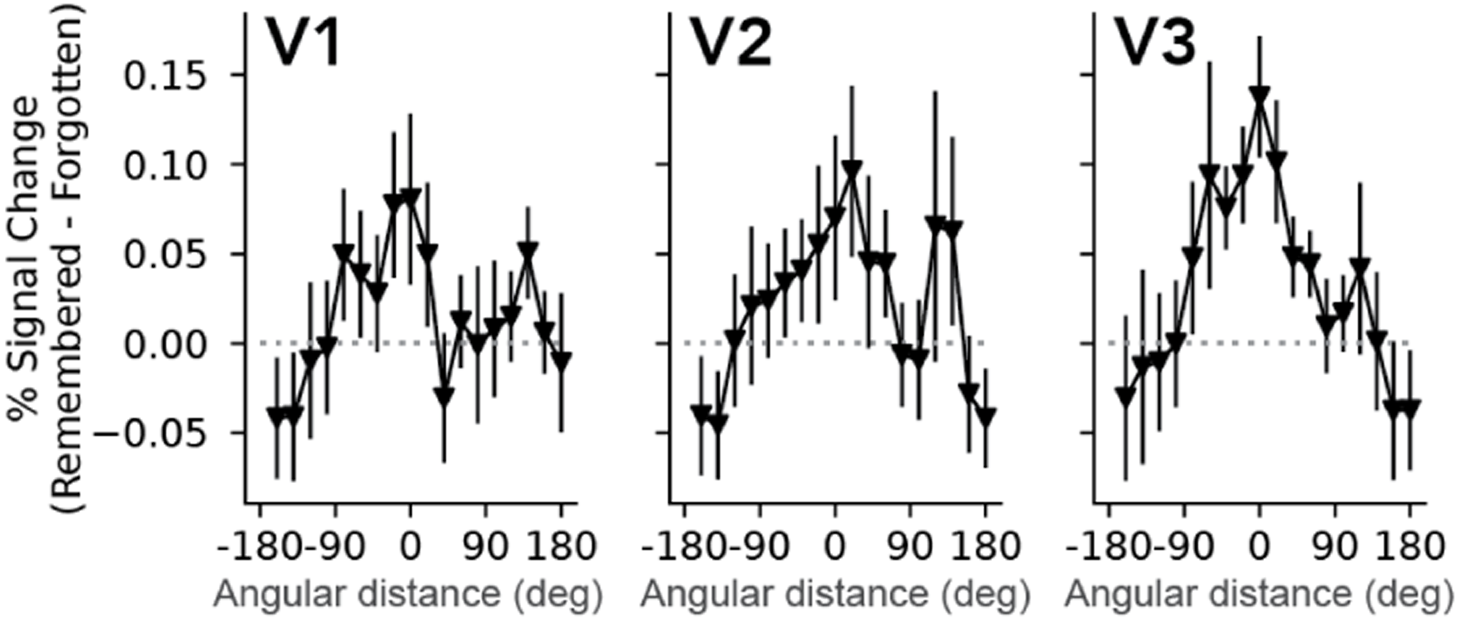
Subsequent memory effect in V1, V2, and V3. Differences in BOLD response between remembered and forgotten trials as a function of angular distance to the target, for each visual map V1, V2, and V3. Error bars indicate one standard deviation of the bootstrapped differences between the two groups. The dashed grey line indicates equality between groups. V3 shows the largest difference between remembered and forgotten responses during encoding, smallest difference in V1.

## Discussion

### Summary

We measured the spatial tuning of visual cortical responses during encoding, recognition, and recall of visually presented objects in a one-shot encoding experiment. We demonstrated that a single encoding experience was sufficient to result in spatially tuned responses in early visual cortex during later memory retrieval. These responses occurred even when spatial information was not explicitly probed, indicating that such responses can arise spontaneously. However, the response amplitude in visual cortex due to memory retrieval was several times lower than what has been found in similar studies with more training prior to retrieval.

Despite the low amplitude of the memory responses, we found multiple relationships between visual cortex activity and memory behavior. First, spatially tuned responses in visual cortex during both recognition and recall were associated with the accuracy of location memory. Second, spatial tuning during encoding – specifically, enhanced responses near the object location and reduced responses elsewhere – was associated with correct responses on the subsequent memory queries. These results point to a strong link between the fidelity of early visual cortical representations and one-shot episodic memory performance. They also demonstrate that the engagement of early sensory areas during memory retrieval does not require highly-trained associations, a common component of lab studies, but rather is a generalizable property of episodic memory, albeit at a low amplitude

### Low amplitude responses during retrieval of one-shot memories

The amplitudes of the one-shot memory responses in visual cortex were lower than in prior studies with highly-trained associations (Favila et al., 2022; Woodry et al., 2025). In those studies, the memory signals were only four times lower than perceptual signals, compared to 25-fold here. The substantial decrease in the relative strength of the memory response here is consistent with the idea that repeated study and retrieval practice increase the strength of the memory response in early visual cortex. Although the amplitude of the V1 to V3 memory responses was low, spatial tuning was evident and was predictive of recall behavior. The relatively low amplitude modulations of early sensory cortex may be more typical of top-down effects than those found when participants are highly practiced at a particular memory.

The encoding model we used facilitated detection and quantification of memory responses in early visual cortex. Indeed, had we not assessed these responses through the lens of an encoding model, we likely would have missed the spatially tuned responses. First, because of low amplitudes and trial-to-trial variability, a typical decoding model would be underpowered. Second, because the cortical population response strongly depends on stimulus location at encoding, univariate analyses that average the V1 cortical response across trials would fail to produce any systematic activation.

### Incidental reactivation in early visual cortex

Strikingly, we observed spatially tuned responses in early visual cortex in the recognition task, when spatial location was not relevant. This indicates that reactivation of perceptual representations can occur in the absence of explicit retrieval demands for spatial information. Why should the brain generate irrelevant signals? One plausible explanation is that retrieval initially reactivates all or many of the features of a memory irrespective of the task demand, after which higher level cortical regions then extract the information relevant for a particular task (Favila et al., 2018; Kuhl et al., 2013; Kwak & Curtis, 2022). Generating only the cortical responses that are task-relevant would require finer and more flexible top-down control. Another possibility is that space is a privileged feature of the memory system, where an object’s location has a high probability of being incidentally encoded. In fact, space has been proposed to be fundamental to episodic memory (O’Keefe & Nadel, 1978) and working memory (Pertzov & Husain, 2014); but see (Woodman, 2021) for a different view).

An alternative possibility is that participants in our experiment knew that they would be tested on object locations, and thus may have been strategically recalling spatial information during the recognition task to practice for the recall task. Spatial tuning in the recognition task, then, could in part be caused by spatial recall. If so, we would expect spatial tuning to increase across the experiment as participants became practiced at this strategy. This was not the case. The fidelity of the spatial response was as strong in the first recognition block — prior to encountering the recall task — as the remainder of the experiment (Supplemental Figure 3-2).

Regardless of whether spatial tuning during recognition reflects strategic covert recall or is spontaneous, our results support the idea that spatial tuning is related to successful memory retrieval and not to other response-related variables. When we aligned memory responses to the reported location for ‘forgotten’ trials, we failed to find tuning present in the recognition and recall responses. This suggests that the spatially tuned memory response in early visual cortex is not caused by the decision (commitment to one of the choices) or the response (button press), but is more likely associated with memory of the location.

### Subsequent memory effects

Interestingly, tuning differences related to successful memory retrieval were evident at encoding: subsequently remembered objects elicited stronger responses near the object’s location and weaker responses elsewhere compared to subsequently forgotten objects. Notably, prior work on subsequent memory effects has not typically implicated early visual cortex (Kim, 2011). Such responses might have been missed since most papers on subsequent memory look for univariate differences across entire regions of interest. Here, we see small, but robust, subsequent memory effects at specific visual field locations on the cortical surface which change on each trial, and which would be obscured by averaging over all of early visual cortex. The spatial specificity of these effects suggests that they are unlikely to be caused by global mechanisms like arousal, which would impact the entire visual field map (Burlingham et al., 2022).

There are several plausible mechanisms that could yield localized gains in the early visual cortex response for subsequently remembered items. First, these effects could be caused by stimulus properties such as image memorability (Rust & Mehrpour, 2020). Image memorability effects differ from subsequent memory effects in that their neural correlates are independent of whether an individual subject remembers the image or not (Bainbridge et al., 2017; Mohsenzadeh et al., 2019). Enhanced responses related to image memorability have thus far only been reported in higher level visual cortex and the medial temporal lobe, and have not been reported in early visual cortex (Bainbridge et al., 2017; Bainbridge & Rissman, 2018). Another candidate mechanism is fluctuations in spatial attention. Attention is known to enhance the neural representation of relevant information in visual cortex (Tootell et al., 1998; Uncapher & Rugg, 2009). Participants likely attended to the location of the target during encoding, and there would naturally be trial-to-trial variation in the level of attention deployed. Selective attention is known to modulate activity in early visual cortex in a similar manner to what we observed (Ling et al., 2015; Tootell et al., 1998; Tünçok et al., 2024) and attention has been repeatedly implicated in promoting memory encoding (Craik et al., 1996; Uncapher & Wagner, 2009; Wager et al., 2004).

### Isn’t it just attention?

The spatially tuned activity we observed in visual cortex during memory retrieval has some similar properties to spatially tuned responses during visual attention tasks (Dugué et al., 2020; Kastner et al., 1998; Tootell et al., 1998; Tünçok et al., 2025). However, there is no visual stimulus at the object’s original location during the memory retrieval tasks, nor an expectation that a stimulus will appear there, nor (in the recognition task) any explicit task related to the location. Therefore, information about the object’s original location must be retrieved from long-term memory storage. Attention can’t explain this retrieval. However, attention might get invoked if the memory is successfully retrieved (Cabeza et al., 2008). This possibility is not specific to spatial tasks but is always present: memory of any visual feature (e.g. color, shape) might similarly invoke feature-based attention.

### Space as an important feature of episodic memory

The visual cortex activity during memory retrieval was related to the location of the stimulus. We chose space as the feature of interest primarily because it is an organizing principle of the visual system (Rodieck, 1998; Wandell, 1995) and because it is a particularly important feature of episodic memory. Indeed brain areas specialized for episodic memory (medial temporal lobe) are also heavily implicated in spatial processing (Hartley et al., 2014; O’Keefe & Nadel, 1978; Ruiz et al., 2020). Lesions to the hippocampus cause a deficit in the ability to retrieve the spatial location associated with an item (Smith & Milner, 1981, 1989). While recognizing the visual features of an object is important, the ultimate utility of episodic memory is to guide future behavior and decision-making (Schacter & Addis, 2007). Because physical actions are inherently spatial, memory for isolated visual features is often insufficient; memory representations must be grounded in spatial frameworks to successfully guide targeted interactions with the environment. Thus, while it is not known whether the same patterns we observed would be found for other visual features during one-shot memory, the fact that it is observed for space is important in and of itself.

### The role of early visual cortex in cognition

Most models of visual cortex have emphasized feedforward processing (Crick & Koch, 1998; Hubel & Wiesel, 1965; Marr, 2010), assigning primary visual areas a limited role in higher-level functions such as memory. An abundance of feedback connections to visual areas are known to exist (Markov et al., 2014), but in the classic view of visual cortex, feedback signals are thought to be modulatory and relatively weak. By contrast, other accounts place a greater emphasis on visual cortex in cognition, proposing that it can serve as a high-resolution buffer or “blackboard” for analyzing images via feedback processes (Gilbert & Sigman, 2007; Lee et al., 1998; Lee & Mumford, 2003; Williams et al., 2008).

Our findings help reconcile these views by showing that early visual cortex is engaged in episodic memory even with minimal demands. We observed memory responses in early visual areas even when task demands did not explicitly encourage imagery or allow for repeated practice, consistent with other evidence for the automatic involvement of sensory systems in internally-oriented cognition (Zokaei et al., 2019). At the same time, these memory signals were approximately 25-fold lower in amplitude than visually evoked responses. The low amplitude may allow early visual cortex to preserve its capacity to faithfully represent incoming sensory input even when reconstructing detailed sensory information during memory retrieval.

## Conclusion

A hallmark of episodic memory retrieval is that it includes sensory details from a single prior experience, contributing to the subjective sense of “mental time-travel” (Tulving, 1985). Yet brain areas such as primary visual cortex, which are best suited to represent sensory details, are not believed to store episodic memories from single experiences; memories are stored in cortex only after repeated exposure or the passage of time (McClelland et al., 1995). We found that episodic memory retrieval from single events engages early visual cortex in a feature-specific manner, and that this activity is correlated with successful memory behavior. Thus, even though memories are not stored in early visual cortex, a core part of the human episodic memory system incorporates early visual cortex in the readout of visual memory and the translation to memory-guided behavior.

## Materials and Methods

### Subjects

20 human subjects (12 females, 25-47 years old) were recruited to participate in the experiment and were compensated for their time. One participant was excluded from the analysis because they misunderstood the instructions for providing behavioral responses during the recall task. An initial group of 5 subjects were used for pilot analyses while the remaining 14 were used to validate those analyses. We report here the analyses across all 19 subjects. Subjects were recruited from the New York University community and included the authors S.E.F. and J.W. Subjects other than the authors were naive to the study’s purpose. Analyses were re-computed after excluding the experimenter subjects to ensure the results hold (see Supplemental Table 1). All subjects gave written informed consent to procedures approved by the New York University Institutional Review Board prior to participation. All subjects had no MRI contraindications, normal color vision, and normal or corrected-to-normal visual acuity.

### Stimuli

#### Object stimuli

This experiment involved testing memory retrieval after a single encoding experience. Therefore, this required the use of a large number of easily recognizable cues with a low probability of confusion. Based on subjective ratings from six subjects not included in the pilot or main experiment, we identified 480 everyday object images from a publicly available dataset that were recognizable in the near periphery (BOSS dataset: (Brodeur et al., 2010). For each set of the three tasks (encoding, recognition, recall), 48 small images of unique objects were sampled without replacement from the 480 images. This produced ten non-overlapping subsets of images — one for each set of three tasks. Within each subset, half (24) were randomly selected without replacement to be used in both the encoding and recall tasks, while the total (48) were used in the recognition task.

On each trial, a small image of one of these objects was shown in either the near periphery (for encoding trials) or at fixation (for recognition and recall trials). During the encoding task, objects were presented at one of four equispaced peripheral locations around the visual field (45°, 135°, 225°, and 315°) at 2° eccentricity from a central fixation point, spanning 2.5° of visual angle. For the recognition and recall tasks, objects were presented at fixation, spanning 1.5° visual angle.

A circular guide spanned the visual field, centered at fixation. This guide consisted of three concentric white circles increasing in radii (2°, 4°, and 6° eccentricity), a horizontal and vertical line that each bisected the circles, and a small white dot at fixation. Object stimuli were presented on top of this guide to provide an external spatial frame of reference.

### Experimental Procedure

The experiment cycled through sets of three scans: encoding, recognition, recall. These sets were repeated five times per scan session (Figure 1). Each subject participated in two scan sessions, for a total of 30 scans (10 per task). Encoding and recall scans consisted of 24 trials, one for each object, in random order. Recognition scans consisted of 48 trials, one for each object from the prior encoding task (old), and another 24 objects not seen before in the experiment (new) — all in random order. Objects in the encoding task were randomly assigned to, and presented in one of, four peripheral locations. Across both scan sessions this resulted in a total of 240 trials each for the encoding and recall tasks, and 480 trials for the recognition task. Behavioral responses on each trial were collected using a four-response button box. Subjects were instructed to maintain central fixation for all tasks. An EyeLink 1000 Plus eye tracker was used to collect gaze position data and ensure fixation. The eye tracker was mounted onto a rig in the magnet bore that holds the projection screen.

For each encoding trial, an object was presented in the periphery for 2 seconds. This was followed by an inter-trial interval lasting 4 to 7 seconds. Subjects were instructed to indicate whether the peripherally presented object was smaller than a shoebox (Button 1), larger than a shoebox (2), or unrecognizable (3).

For each recognition trial, an old (from the prior encoding task) or new object was presented at fixation for 2 seconds. This was followed by an inter-trial interval lasting 4-7 seconds. Subjects were asked to indicate whether the centrally presented object was old (1), or new (2).

For each recall trial, an old object (from the prior encoding task) was presented at fixation for 2 seconds. This was followed by an inter-trial interval lasting 4-7 seconds. Subjects were asked to indicate the object’s original location from the prior encoding task (1: upper right, 2: lower right, 3: lower left, 4: upper left).

Note that while the objects encoded were unique, the 4 locations probed were not. Thus In our task, there was a many-to-one mapping between the cue (trial unique object) and contextual feature (one of four spatial locations), such that each object-location pairing was only encoded once, but locations repeated over trials. This has a strong precedent in the episodic memory literature, where similarly structured contextual or source recollection tasks have been used for decades. These tasks frequently have non-unique contextual features, but are hippocampally-mediated (Cooper & Ritchey, 2020; Mugikura et al., 2010; Uncapher et al., 2006; S. S. Yu et al., 2012), suggesting that they are episodic memory tasks. Indeed, most everyday episodic memories, such as remembering what you ate for breakfast, contain non-unique features.

#### Retinotopic mapping procedure

Each participant additionally underwent 10-12 retinotopic mapping scans in a separate scan session. The stimuli and procedures for the retinotopic mapping session followed those used in Himmelberg et al. (2021), summarized briefly here. Each scan involved contrast patterns presented within bar apertures (1.5° width), which swept across the visual field within a 12°-radius circle. There were 8 bar sweep types in total, comprising 4 diagonal directions and 4 cardinal directions. Each bar sweep took 24 seconds to complete: cardinal sweeps took 24 seconds to traverse the full extent of the circular aperture; diagonal sweeps took 12 seconds to move to the halfway point, subsequently followed by 12 second long blank periods. Each functional scan comprised 8 sweeps, for a total of 192 seconds. The contrast patterns featured pink noise (grayscale) backgrounds with randomly sized and placed items, updated at 3 Hz. Participants were instructed to press a button whenever the fixation dot changed color, which occurred approximately once every three seconds. These contrast patterns were originally implemented by Benson et al. (2018).

### MRI Acquisition

A 3T Siemens Prisma MRI system and a Siemens 64-channel head/neck coil were used to collect functional and anatomical images at the Center for Brain Imaging at New York University. We obtained functional images using a T2*-weighted multiband echo planar imaging (EPI) sequence with whole-brain coverage (repetition time = 1 s, echo time = 37 ms, flip angle = 68°, 66 slices, 2 x 2 x 2 mm voxels, multiband acceleration factor = 6, phase-encoding = posterior-anterior), developed at the University of Minnesota’s Center for Magnetic Resonance Research (Feinberg et al., 2010; Moeller et al., 2010; Xu et al., 2013). To estimate and correct susceptibility-induced distortions in the functional EPIs, we collected spin echo images with both anterior-posterior and posterior-anterior phase-encoding. Additionally, for each of the 20 subjects, we acquired one to three whole-brain T1-weighted MPRAGE 3D anatomical volumes (.8 x .8 x .8 mm voxels).

### MRI Processing

We used *pydeface* (https://github.com/poldracklab/pydeface) to deface and anonymize all original MRI data (DICOM files). We then used the Heuristic Dicom Converter (Halchenko et al., 2018) to convert this data to NIFTI and organized the files into the Brain Imaging Data Structure format (K. J. Gorgolewski et al., 2016). We then preprocessed the data using fMRIPrep 20.2.7 (Esteban et al., 2018, 2019), which is based on Nipype 1.7.0 (K. Gorgolewski et al., 2011; K. J. Gorgolewski et al., 2018).

The following sections on anatomical and functional data preprocessing are provided by the fMRIPrep boilerplate text generated by the preprocessed scan output.

#### Anatomical data preprocessing

Each of the one to three T1w images was corrected for intensity non-uniformity with N4BiasFieldCorrection (Tustison et al., 2010), distributed with ANTs 2.3.3 (Avants et al., 2008). The T1w-reference was then skull-stripped with a Nipype implementation of the antsBrainExtraction.sh workflow (from ANTs), using OASIS30ANTs as target template. Brain tissue segmentation of cerebrospinal fluid, white-matter and gray-matter was performed on the brain-extracted T1w using fast (FSL 5.0.9, (Zhang et al., 2001)). A T1w-reference map was computed after registration of the T1w images (after intensity non-uniformity-correction) using mri_robust_template (FreeSurfer 6.0.1, (Reuter et al., 2010)). Brain surfaces were reconstructed using recon-all (FreeSurfer 6.0.1, (Dale et al., 1999)), and the brain mask estimated previously was refined with a custom variation of the method to reconcile ANTs-derived and FreeSurfer-derived segmentations of the cortical gray-matter of Mindboggle (Klein, 2017).

#### Functional data preprocessing

For each of the 30 BOLD runs found per subject (across all tasks and sessions), the following preprocessing was performed. First, a reference volume and its skull-stripped version were generated by aligning and averaging a single-band reference. A B0-nonuniformity map (or fieldmap) was estimated based on two EPI references with opposing phase-encoding directions, with 3dQwarp (Cox & Hyde, 1997). Based on the estimated susceptibility distortion, a corrected EPI reference was calculated for a more accurate co-registration with the anatomical reference. The BOLD reference was then co-registered to the T1w reference using bbregister (FreeSurfer) which implements boundary-based registration (Greve & Fischl, 2009). Co-registration was configured with six degrees of freedom. Head-motion parameters with respect to the BOLD reference (transformation matrices, and six corresponding rotation and translation parameters) are estimated before any spatiotemporal filtering using mcflirt (FSL 5.0.9, (Jenkinson et al., 2002)). BOLD runs were slice-time corrected to 0.445s (0.5 of slice acquisition range 0s-0.89s) using 3dTshift from AFNI 20160207 (Cox & Hyde, 1997). First, a reference volume and its skull-stripped version were generated using a custom methodology of fMRIPrep. The BOLD time-series were resampled onto the fsnative surface. The BOLD time-series (including slice-timing correction) were resampled onto their original, native space by applying a single, composite transform to correct for head-motion and susceptibility distortions. These resampled BOLD time-series will be referred to as preprocessed BOLD. All resamplings can be performed with a single interpolation step by composing all the pertinent transformations (i.e. head-motion transform matrices, susceptibility distortion correction, and co-registrations to anatomical and output spaces). Gridded (volumetric) resamplings were performed using antsApplyTransforms (ANTs), configured with Lanczos interpolation to minimize the smoothing effects of other kernels (Lanczos, 1964). Non-gridded (surface) resamplings were performed using mri_vol2surf (FreeSurfer).

Many internal operations of fMRIPrep use Nilearn 0.6.2 (Abraham et al., 2014), mostly within the functional processing workflow. For more details of the pipeline, see the section corresponding to workflows in fMRIPrep’s documentation.

#### General linear models

From each subject’s surface based time series, we used GLMSingle (Prince et al., 2022) to estimate the neural pattern of activity during the 2 second stimulus presentations for each trial. GLMsingle first fits to each vertex’s time series an optimal response function from a bank of 20 candidate hemodynamic response functions (Natural Scenes Dataset, (Allen et al., 2021)). Second, noisy vertices are identified from the data (defined by negative *R^2^*) and then used to estimate noise regressors. This step iteratively uses principal component analysis to derive the noise regressors, where an optimal number of noise regressors is chosen and then projected out of the time series’ data. Third, fractional ridge regression is used to improve the robustness of single-trial beta estimates, particularly useful here as our design relies on single-trial encoding, recognition, and retrieval of object stimuli.

We constructed three separate design matrices to model the BOLD time series using GLMSingle, one for each of the three tasks. Each of the three design matrices has a regressor corresponding to the object’s true location (‘target-aligned’). Recognition design matrices had an additional regressor for ‘new’ objects.

#### Population receptive field models

Data from the retinotopy session were used to fit a population receptive (pRF) model to each vertex on the cortical surface, as described by Himmelberg et al. (2021); section 2.6). To briefly summarize, we fit a circular 2D-Gaussian linear pRF to each surface vertex’s BOLD time series, averaged across identical runs of the bar stimulus. These fits were implemented with *Vistasoft* software (Dumoulin & Wandell, 2008), in conjunction with a wrapper function to handle surface data (https://github.com/WinawerLab/prfVista). Gaussian center (*x, y*) and standard deviation (σ) comprised the pRF model parameters.

### Visual field map definitions

We used the visual tool *cortex-annotate* (https://github.com/noahbenson/cortex-annotate), which is built on *neuropythy* software (Benson & Winawer, 2018), to draw and define visual field maps for each subject. To do so, we traced the polar angle reversals on each subject’s cortical surface. We followed common heuristics to define four maps spanning early visual cortex: V1, V2, V3 (Benson et al., 2022; Himmelberg et al., 2021). We then defined experiment-specific regions of interest for each visual field map. To do this we excluded vertices whose variance explained was less than 10% and whose pRF centers were outside one σ of 2° (the target eccentricity in the experiments). For example, a vertex with a pRF center at 1° and pRF size (σ) of 0.5° would be excluded, but a vertex with pRF center at 3° and pRF size of 1.5° deg would be included. We restricted vertices in our analyses this way to examine polar angle activation profiles, described in the next section. These mapping estimates are solely based on the retinotopy scans, and are therefore independent of the main experiment.

### Quantifying behavioral performance at recognition and recall

#### Object recognition

During the recognition task, subjects responded with either ‘new’ or ‘old’ for objects presented at fixation. Half of these objects were present in the previous encoding task (‘old’), the rest ‘new’. Hits were classified as trials where previously seen objects were correctly identified as ‘old’. Misses were classified as trials where previously seen objects were incorrectly identified as ‘new’. Correct rejections were classified as trials where new objects were correctly identified as ‘new’. False alarms were classified as trials where new objects were incorrectly identified as ‘old’. For each scan, the total number of hits were divided by the total number of ‘old’ objects, to get hit rate as a percentage. Likewise, the total number of false alarms was divided by the total number of ‘new’ objects to get the false alarm rate as a percentage. Miss and correct rejection rates were calculated as the complement of the hit and false alarm rates, respectively.

#### Spatial location recall

During the recall task, subjects reported which of four locations the object presented at fixation originally appeared at (from the encoding task). Trials were classified as accurate or inaccurate if the reported location matched that of the original location, or source location, of the object. Therefore, because all objects shown were ‘old’ objects that had appeared within the encoding task in one of four locations, the chance level was at 25%. For each scan, the total number of accurate responses were divided by the number of trials to get mean accuracy.

### Gaze position analysis

The gaze position of the participants’ right eye (in X, Y screen coordinates) was recorded at a sampling rate of 1 kHz while undergoing the three tasks in the MRI scanner. Five subjects were excluded from this analysis due to lack of eyetracking data (including the two experimenter subjects), one other subject was excluded because of persistent failure to calibrate the eyetracker during the two sessions. Gaze position data was converted from X and Y positions in screen pixel coordinates to degrees in visual angle. We removed blink epochs identified by the Eyelink software using *pynapple* software (Viejo et al., 2023). Within each run, we corrected for calibration errors by subtracting the median gaze position during the 300ms before stimulus onset from the gaze measured during stimulus display. For each trial we averaged the X, Y position across time points during the two second stimulus display. We then averaged these time-averaged values across all trials with the same stimulus source location, producing four X-Y pairs per participant and each of the three tasks.

### Analyses quantifying spatial tuning in visual cortex at encoding and retrieval

We computed polar-angle activation profiles for each task (encoding, recognition, recall) across a region of interest comprising visual maps V1, V2, and V3. To do so, we first obtained response amplitudes from the outputs of GLMsingle for each vertex on the cortical surface and each trial. PRF mapping, which was conducted in a separate fMRI session, returned visual field coordinates for each vertex. We excluded vertices not included in the ROI definitions outlined above. We further restricted trials by several criteria: for all tasks, we only included trials where behavioral responses were provided. In the recognition task, we only included trials that were classified as ‘hits’ (subject reported ‘old’ when viewing an ‘old’ object). Some analyses further restricted trials across all tasks to those whose object locations were accurately reported in the recall task. We binned the response amplitudes for the remaining trials and vertices by the angular distance between each vertex’s preferred polar angle and the source location for that object. This binning by angular distance allows for averaging trials across the four peripheral source locations. This produces an activation profile as a function of distance from the object’s source location; an activation profile for each subject and task. Each activation profile is then normalized by dividing by the vector length of the perceptually evoked activation profile (from the encoding task) for the corresponding subject. We then preserved meaningful units by rescaling each activation profile by the average vector lengths from the encoding task. Each of the resulting averaged activation profiles peaked near 0°, where values decreased moving away from the peak response. We therefore fit a von Mises distribution to the averaged activations as a function of angular distance to the source location, θ_source_ (Eq. 1). From each of these fits we estimated the peak location, the amplitude (peak to trough), and the width (full-width-at-half-maximum; FWHM).

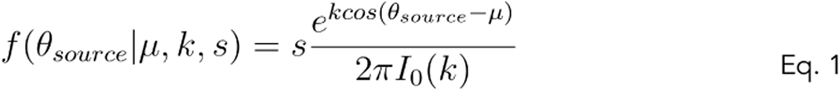

Each metric provides a quantitative measure of some aspect of the tuning. However, on their own they can not provide a holistic picture of the quality of the tuning; for example, the response may be narrow but peak at a location opposite the source. Therefore to summarize the degree to which the tuned response concentrates at the source location we computed an angular “fidelity” metric which normalizes the length of the vector sum of the polar angle activation profile projected to 0° — a modified version of the circular fidelity metric used in prior related work (Vo et al., 2022).

To compute this metric, we first baseline subtracted each polar angle activation profile to remove any negative values. We then defined each binned activation (w_j_) to be a vector pointing in the direction of the bin location (ɑ_j_) with a length equal to the activation, and then summed all 18 component vectors. We normalized the resultant vector by dividing the summed length of all component vectors. The direction of the normalized resultant vector captures the circular mean of each activation profile (Eq. 2), and the resultant vector length captures how concentrated the activations are near the mean direction (Eq. 3). Both the circular mean and resultant vector length are metrics derived from the CircStats MATLAB toolbox (Berens, 2009). The fidelity metric is obtained by multiplying the cosine of the circular mean, which captures how far away the mean is from 0°, by the resultant vector length (Eq. 4). Because cosine goes from -1 to 1, and the resultant vector length is normalized to be between 0 and 1, the fidelity metric is bound between +1 and -1. A value of 1 indicates a BOLD amplitude of 0 in every bin except the one centered at 0° (maximum directional bias towards 0°), a value of -1 indicates an amplitude of 0 everywhere except the bin at 180° (maximum directional bias away from 0°), and a value of 0 indicates no directional bias.

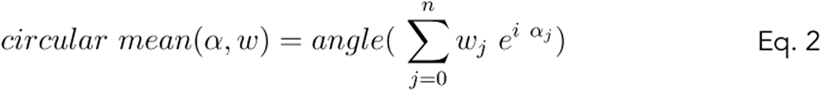

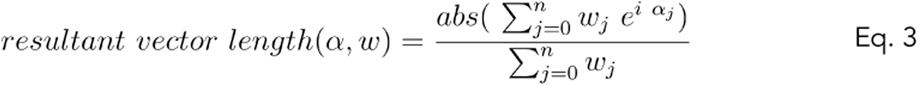

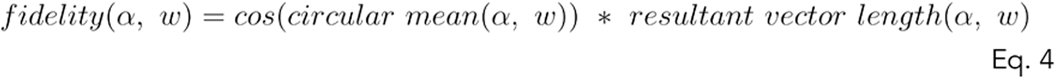

We repeated the von Mises and fidelity analyses, bootstrapping across subjects with replacement 10,000 times to obtain 68% and 95% confidence intervals for each of these four tuning metrics.

To assess the quality of tuning at the participant level, we computed the null distribution separately for each participant by scrambling the assignment of target locations across trials, and then re-computing the fidelity. This was done 1,000 times per participant.

#### Resampling statistics

We analyzed resampled data to infer spatial tuning properties as a function of condition and of other trial-level factors. Statistics reported in this paper are derived from these resampled analyses. We report the mean and confidence intervals (CI) obtained from these analyses to assess our main claims. We report the 95% CI (corresponding to ±2 standard deviations of a normal distribution) when drawing comparisons between a measurement and a fixed value. When comparing two estimates, we instead report the 68% CIs (corr. ±1 standard deviation).

### Software

Model fitting, data visualization, and statistical quantification for all analyses described in this paper were made using matplotlib 3.5.2 (Hunter, 2007), nibabel 3.2.2 (Brett et al., 2022), pandas 1.4.2 (The pandas development team, 2024), scikit-learn 1.0.2 (Pedregosa et al., 2011), scipy 1.8.0 (Virtanen et al., 2020), and seaborn 0.11.2 (Waskom, 2021). Functions to compute circular statistics were converted to python from the CircStats MATLAB toolbox (Berens, 2009). Pynapple was used to process eyetracking data (Viejo et al., 2023).

### Data Availability

The data and code used to reproduce the analyses in this paper can be found at https://osf.io/a9hkg/.

## Acknowledgements

We thank New York University’s Center for Brain Imaging for technical support. This research was supported by the National Institutes of Health (F99 NS105223 to S. Favila; R01 EY027401 to J. Winawer; P30 EY013079 Core grant for vision; R01 MH111417 to J. Winawer; T90 DA059110 computational neuroscience training grant supporting R. Woodry; T32 EY007136 vision neuroscience training grant supporting R. Woodry) and pilot grants from the NYU Center for Brain Imaging. This work was supported in part through the NYU IT High Performance Computing resources, services, and staff expertise.

---

**Supplemental Table 1.** Reanalysis excluding experimenter subjects.

| *Analysis* | *Including experimenter subjects* | *Excluding experimenter subjects* |
| --- | --- | --- |
| **Memory performance for object recognition and spatial location recall** |  |  |
| Average recognition hit rate | 90% [95 CI 87%, 92%] | 90% [87%, 92%] |
| Average recognition false alarm rate | 7% [95 CI 5%, 9%] | 7% [5%, 10%] |
| Average recall accuracy | 69% [95 CI 63%, 75%] | 69% [63% 74%] |
| **A single viewing is sufficient for spatially tuned memory responses in early visual cortex** |  |  |
| Peak location absolute error: encoding | 1° [95 CI 0°, 2°] | 1° [0°, 2°] |
| Peak location absolute error: recognition | 8° [95 CI 0°, 31°] | 9° [0°, 46°] |
| Peak location absolute error: recall | 13° [95 CI 0°, 48°] | 18° [1, 70°] |
| Peak location absolute error: recognition - encoding | 7° [95 CI -1°, 32°] | 8° [-1°, 66°] |
| Peak location absolute error: recall - encoding | 12° [95 CI -1° 55°] | 17° [0°, 82°] |
| Amplitude: encoding - recognition | 0.86% [95 CI 0.69%, 1.03%] | 0.9% [0.71%, 1.08%] |
| Amplitude: encoding - recall | 0.87% [95 CI 0.7%, 1.03%] | 0.91% [0.7%, 1.07%] |
| FWHM: encoding | 81° [68 CI 79°, 83°] | 80° [78°, 81°] |
| FWHM: recognition | 122° [68 CI 97°, 136°] | 115° [71°, 134°] |
| FWHM: recall | 161° [68 CI 111°, 177°] | 150° [91°, 172°] |
| Fidelity: encoding | 0.67 [95 CI 0.62, 0.71] | 0.67 [0.63, 0.71] |
| Fidelity: recognition | 0.38 [95 CI 0.11, 0.49] | 0.36 [0.04, 0.47] |
| Fidelity: recall | 0.32 [95 CI 0.07, 0.46] | 0.26 [0.01, 0.39] |
| **Memory responses in early visual cortex track locations of remembered objects** |  |  |
| *Source-aligned* |  |  |
| Fidelity: recognition remembered | 0.35 [95 CI 0.13, 0.44] | 0.29 [0.08, 0.42] |
| Fidelity: recognition forgotten | 0.24 [95 CI -0.08, 0.34] | 0.24 [-0.16, 0.34] |
| Fidelity: recall remembered | 0.35 [95 CI 0.09, 0.52] | 0.27 [0.05, 0.46] |
| Fidelity: recall forgotten | -0.03 [95 CI -0.14, 0.18] | -0.03 [-0.16, 0.21] |
| *Report-aligned* |  |  |
| Fidelity: recognition forgotten | 0.25 [95 CI -0.13, 0.36] | 0.20 [-0.17, 0.35] |
| Fidelity: recall forgotten | 0.11 [95 CI -0.21, 0.30] | 0.05 [-0.26, 0.24] |
| **Spatial tuning at encoding predicts subsequent memory** |  |  |
| Amplitude: encoding remembered - forgotten | 0.11% [95 CI 0.01%, 0.22%] | 0.09% [-0.02%, 0.21%] |

